# Limitation of alignment-free tools in total RNA-seq quantification

**DOI:** 10.1101/246967

**Authors:** Douglas C. Wu, Jun Yao, Kevin S. Ho, Alan M. Lambowitz, Claus O. Wilke

## Abstract

**Background:** Alignment-free RNA quantification tools have significantly increased the speed of RNA-seq analysis. However, it is unclear whether these state-of-the-art RNA-seq analysis pipelines can quantify small RNAs as accurately as they do with long RNAs in the context of total RNA quantification.

**Result:** We comprehensively tested and compared four RNA-seq pipelines on the accuracies of gene quantification and fold-change estimation on a novel total RNA benchmarking dataset, in which small non-coding RNAs are highly represented along with other long RNAs. The four RNA-seq pipelines were of two commonly-used alignment-free pipelines and two variants of alignment-based pipelines. We found that all pipelines showed high accuracies for quantifying the expressions of long and highly-abundant genes. However, alignment-free pipelines showed systematically poorer performances in quantifying lowly-abundant and small RNAs.

**Conclusion:** We have shown that alignment-free and traditional alignment-based quantification methods performed similarly for common gene targets, such as protein-coding genes. However, we identified a potential pitfall in analyzing and quantifying lowly-expressed genes and small RNAs with alignment-free pipelines, especially when these small RNAs contain mutations.

## Background

RNA-seq continues to pose great computational and statistical challenges. These challenges range from accurately aligning sequencing reads to accurate inference of gene expression levels [1, 2]. The central computational problem in RNA-seq remains the efficient and accurate assignment of short sequencing reads to the transcripts they originated from and using this information to infer gene expressions [3–6]. Conventionally, read assignment is carried out by aligning sequencing reads to a reference genome, such that relative gene expressions can be inferred by the alignments at annotated gene loci [2, 7]. These alignment-based methods are conceptually simple, but the read-alignment step can be time-consuming and computationally intensive despite recent advances in fast read aligners [4, 8, 9]. Recently, several novel tools introduced alignment-free transcript quantification utilizing k-mer-based counting algorithms [4–6]. These alignment-free pipelines are orders of magnitude faster than alignment-based pipelines, and they work by breaking sequencing reads into k-mers and then performing fast matches to pre-indexed transcript databases [4]. To achieve fast transcript quantification without compromising quantification accuracy, different sophisticated algorithms were implemented in addition to k-mer counting, such as pseudoalignments (Kallisto [5]) or quasi-mapping along with GC and sequence-bias corrections (Salmon [6]). Given the wide variety of choices in RNA-seq tools, several studies have benchmarked subsets of read aligners and quantification software. These studies generally suggest that most of the current RNA-seq tools are comparably accurate [10–12].

However, the existing benchmarking studies were generally carried out on either simulated RNA-seq datasets [12] or RNA-seq datasets that focused only on long RNAs, such as messenger RNAs (mRNAs) and long non-coding RNAs (lncRNAs) [10, 11, 13, 14]. Consequently, they did not evaluate whether these tools are suitable for total RNA quantification in datasets that include small RNAs, such as transfer RNAs (tRNAs) and small nucleolar RNAs (snoRNAs). To some extent, the lack of a comprehensive comparison between small and long RNA quantification may be due to the inability of most current RNA-seq methods to efficiently recover these small RNAs [15]. Recently, however, a novel method has overcome this problem by using a thermostable group II intron reverse transcriptase (TGIRT) during RNA-seq library construction [15]. This method enables more comprehensive profiling of full-length structured small non-coding RNAs (sncRNA) along with long RNAs in a single RNA-seq library workflow [15–17]. Thus, it is now possible to benchmark RNA-seq quantification tools on structured small non-coding RNAs.

To address whether current RNA-seq tools can quantify small RNAs as accurately as they do with long RNAs, we tested four gene quantification pipelines on a previously sequenced TGIRT RNA-seq (TGIRT-seq) dataset [15] obtained from the well-studied microarray/sequencing quality control consortium (MAQC) sample set [18, 19]. Of the four tested pipelines, two are alignment-based and two are alignment-free. We found that all four pipelines are mostly concordant in quantifying common differentially-expressed gene targets, such as mRNAs and mRNA-like spike-ins. However, with respect to quantifying small or lowly-expressed genes, we found that the alignment-based pipelines significantly outperformed the alignment-free pipelines.

## Results

### Study design

We tested four RNA-seq quantification pipelines, including two alignment-free and two alignment-based pipelines (Fig. 1): (A) Kallisto, a k-mer counting software that uses pseudoalignments for reducing quantification error and improving speed [5]; (B) Salmon, another k-mer counting software that learns and corrects sequence-specific and GC biases on-the-fly, in addition to using quasi-mapping for further improvement in transcript quantification [6]; (C) HISAT2+featureCounts, a conventional alignment-based pipeline aligning sequencing reads to human genome by a splice-aware aligner, HISAT2 [9], and quantifying genes by featureCounts [3]; and (D) TGIRT-map, a customized alignment-based pipeline using an iterative genome-mapping procedure (Additional File 1).

**Figure 1.**
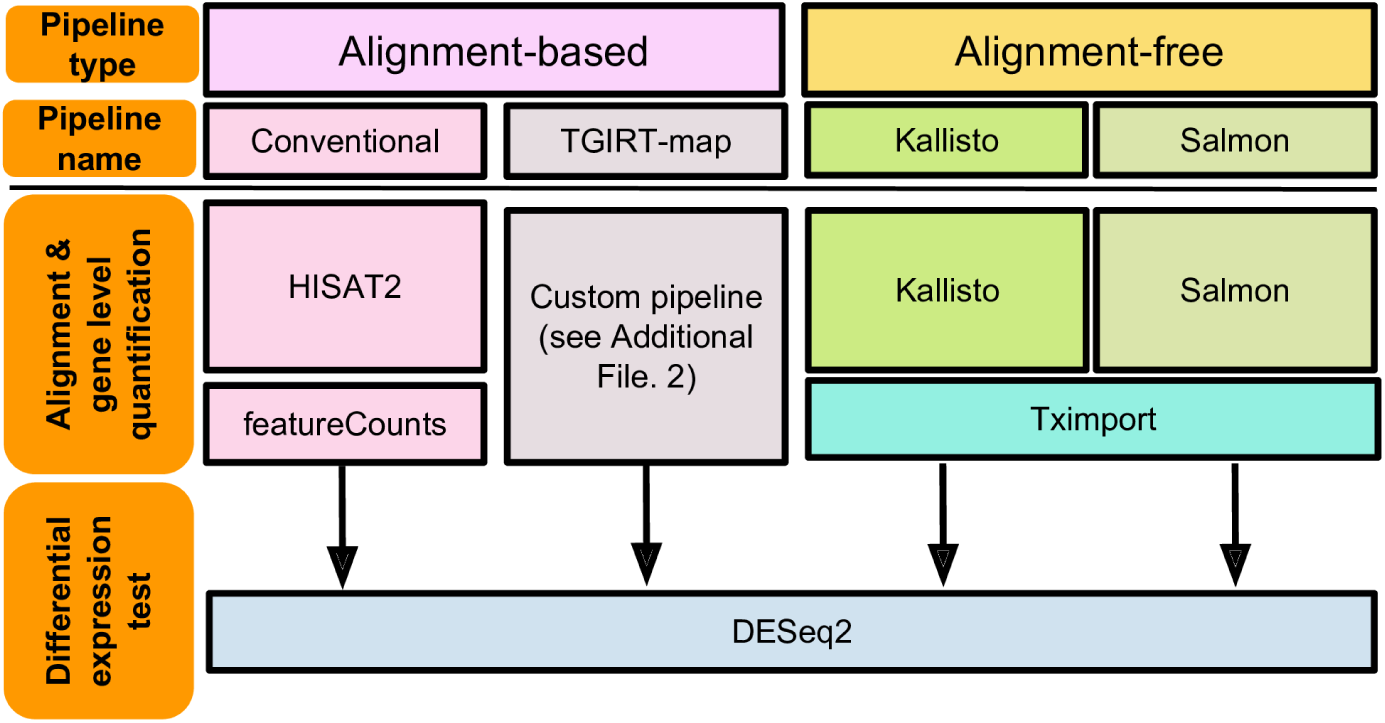
Analysis pipelines and experimental design. We used two pipelines each for the alignment-based and alignment-free approach. The alignment-based pipelines consisted of (A) a HISAT2+featureCounts pipeline using HISAT2 [9] for aligning reads to the human genome and using featureCounts [3] for gene counting, and (B) TGIRT-map, a customized pipeline for analyzing TGIRT-seq data. Further details regarding the custom TGIRT-map pipeline are provided in Methods and in Additional File 1. Two alignment-free tools, Kallisto [5] and Salmon [6], were used for quantifying transcripts. For alignment-free tools, gene-level abundances were summarized by T×import [35]. All differentially-expressed gene callings were done by DESeq2 [34].

The benchmarking dataset we used here consists of TGIRT-seq libraries for four well-defined samples (samples A-D) from the microarray/sequencing quality control consortium (MAQC [18, 19]), each obtained in triplicate [15]. The MAQC samples A and B represent universal human reference total RNA and human brain reference total RNA, respectively, that are mixed with corresponding External RNA Controls Consortium (ERCC) spike-in transcripts. Samples C and D are mixtures of samples A and B at different ratios [18, 19]. This known mix ratios in samples C and D allow for the calculation of expected fold-changes between samples C and D from the measured fold-changes between samples A and B for each gene [18, 19]. Mapping statistics of all pipelines are summarized in Additional File 2 Supplementary Tables S1-4.

### Gene detection and quantification

To test if all four pipelines produced concordant gene quantifications, we first compared the numbers of detected genes across all methods. We considered a gene as detected if it was assigned with a transcripts per million (TPM) value > 0.1. We found that the numbers of detected genes were similar among all tested pipelines (Fig. 2a). Moreover, by comparing the identities of the detected genes, we found that the vast majority of the genes were detected by all tested pipelines (Additional File 3). However, we also found that Salmon and TGIRT-map consistently detected more genes compared to the other two pipelines (Friedman test, *p* = 4×10^−7^; Paired Wilcoxon test, *p* = 0.57 for Salmon vs TGIRT-map, *p* = 5×10^−4^ for other pipeline comparisons).

**Figure 2.**
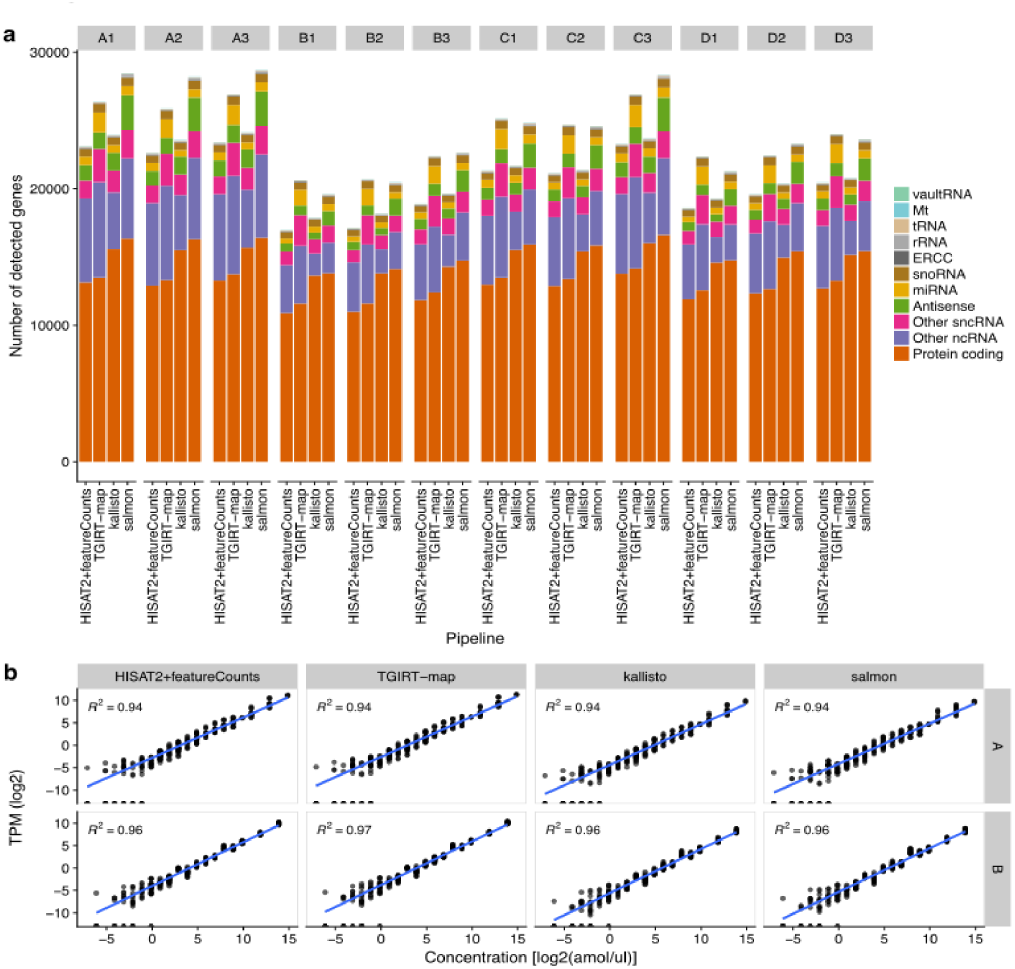
Gene detection and quantification. (a) Numbers and types of detected genes in every sample and pipeline. Genes with TPM > 0.1 were labeled as being detected. The stacked bar charts indicate the numbers of genes detected by each pipeline. The bar charts are grouped by library (A1–D3), where A–D represent the RNA samples and the numbers represent replicate identi ers. The stacked-bars are color-coded by RNA type. (b) ERCC spike-in quantifications versus the true spike-in abundances. Log2 transcripts per million (TPM) values for every ERCC transcript from every replicate are plotted against the known spike-in concentrations, grouped by pipelines and samples. Blue lines indicate least-square regression lines. Coefficients of determination are annotated in each panel.

Even though Salmon and TGIRT-map both detected more genes than did the other two pipelines, the additional genes detected were different. Salmon primarily recovered more long RNAs (labeled as antisense, other ncRNAs, and protein-coding genes; Additional File 3). This enrichment in long RNAs could be pipeline-type-specific (when compared to alignment-based pipelines) or algorithm-specific (when compared to Kallisto). Pipeline-type-specific differences could be due to the probabilistic gene quantification methods of Salmon [6]. While Salmon can assign fragments to multiple genes, each fragment can only be assigned to a single gene by the alignment-based pipelines under our parameters. In terms of algorithm, the result might be due to how Salmon corrects for GC and sequencing biases or how it handles equivalent classes (i.e. multiply-mapped reads) relative to Kallisto [6]. On the other hand, TGIRT-map recovered more miRNAs (likely to be mis-counting of fragmented exons or unannotated exons in these libraries), some long non-coding RNAs (annotated as other ncRNAs), and small non-coding RNAs (annotated as other sncRNA) (Additional File 3). These enrichments under TGIRT-map could be pipeline-type-specific when compared to alignment-free pipelines, which may be affected by the choice of k-mer size. The differences between TGIRT-map and HISAT2+featureCounts were possibly the result of an additional alignment step (BOWTIE2) [20] after the spliced-read mapping step (HISAT2) [9] in TGIRT-map (Additional File 1).

To evaluate if gene expression level estimates were concordant among the tested pipelines, we made pairwise comparisons of these estimates between pipelines (Additional File 4). The gene expression level estimates were generally highly correlated, with Pearson’s correlations ranging from 0.67-0.99 (Additional File 4). The Pearson’s correlation coefficients were consistently very high for pairwise comparisons between alignment-free pipelines (Kallisto vs Salmon; 0.98-0.99) or between alignment-based pipelines (HISAT2+featureCounts vs TGIRT-map; 0.94-0.95). By contrast, any pairwise correlation between an alignment-free tool and an alignment-based pipeline was generally lower (0.67-0.72; Additional File 4).

Further analyses revealed that different gene types showed distinct variations in gene expression level correlations for every pairwise pipeline comparison (Additional File 4). For instance, ERCC spike-ins, *in vitro* transcripts that mimic protein-coding transcripts, were recovered with very high correlations for all pairwise pipeline comparisons (Pearson’s correlations: 0.99-1; Additional File 4). In comparison to the true abundances that were spiked into the RNA samples [15], the relative expression levels of these ERCC spike-ins were estimated as they were designed and tightly correlated to their true concentrations. We observed a near-perfect linear relationship between inferred TPM values and true concentrations (both log-transformed) for all pipelines (Fig. 2b; *R*^2^ > 0.94 for every sample and pipeline; Kruskal-Wallis-test *p* = 0.346). By contrast to ERCC transcripts, the gene expression level estimates of other common gene targets (antisense, protein-coding genes, etc.) were not as highly correlated among tested pipelines (Additional File 4). To identify the source of this discrepancy, we divided genes into quantile groups of gene lengths or gene expression levels, and found that the abundance estimation inconsistencies among pipelines were largely caused by short gene lengths and low expression levels (Additional File 5), as suggested previously [13].

### Differential expression measurements of long genes

The most popular application of RNA-seq is the detection of differentially-expressed genes. To compare the accuracy of differential expression inferences among each pipeline, we plotted the deviation of measured log2 fold-changes to the expected log2 fold-changes between samples A and B for every ERCC spike-in transcript (Fig 3a; 23 ERCC transcripts in each expected differentially-expressed group). For all pipelines, fold-changes between samples A and B were mostly underestimated (negative Δlog2 fold-changes; Fig. 3a). To quantify the accuracies of fold-change detections for each method, we computed root mean square errors (RMSE) for each ERCC group (Additional File 2). Comparisons of the RMSE values for each ERCC group among pipelines indicated that alignment-free pipelines had comparable performances to alignment-based pipelines in estimating differential expression of ERCC spike-ins (Friedman-test *p* = 0.016; two-sided paired Wilcoxon-test *p* > 0.1 for all pairwise comparisons).

**Figure 3.**
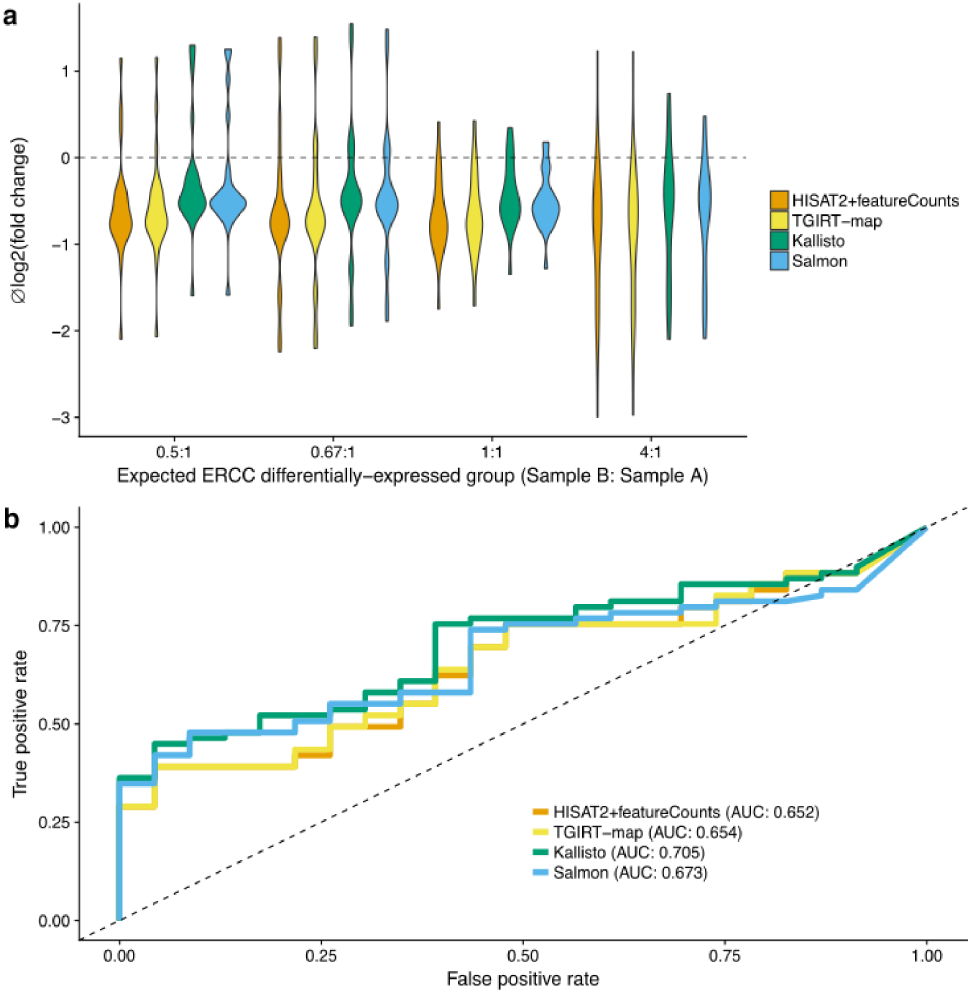
Differential expression analysis of ERCC spike-ins. (a) Violin-plots of deviations between measured and expected log2 fold-changes of ERCC transcript between samples A and B. The distributions of log2 fold-change errors for each ERCC transcript are grouped by their expected differentially-expressed groups and are color-coded by the tested pipelines. The horizontal dashed line indicates no error. (b) Receiver operating characteristic curves for calling ERCC spike-ins as differentially-expressed. Areas under the curve were computed using p-values assigned by the differential expression caller (DESeq2 [34]) on abundance estimations of each ERCC transcript from each pipeline.

To test whether the four pipelines provided reliable gene expression level estimates for calling differentially-expressed genes, we plotted the receiver operating characteristic (ROC) curves for the prediction of ERCC spike-in transcripts differentially-expressed status. For every pipeline, we used the *p*-values assigned to the differential expression tests for binary prediction cutoffs and the known spike-in concentrations for assigning true differential-expression status (Fig. 3b). By design, there are 23 spike-ins with same concentrations and 69 spike-ins with different concentrations between samples A and B [21]. Using area under the curves (AUC) of the ROC curves, we found that alignment-free tools (Kallisto and Salmon) performed slightly better than alignment-based pipelines in accurately calling differentially-expressed spike-in transcripts (AUC: 0.65, 0.65, 0.71, and 0.68 for HISAT2+featureCounts, TGIRT-map, Kallisto, and Salmon respectively; one non-differentially-expressed and two differentially-expressed ERCC spike-ins had TPM = 0 in all pipelines). In addition to analyzing *in vitro* ERCC transcripts, we also verified that all pipelines performed nearly identically in terms of quantifying *in vivo* transcripts, by comparing estimated expression levels to TaqMan qRT-PCR results published previously (Additional File 6).

### Whole transcriptome differential expression analysis

To benchmark the suitability of different gene quantification pipelines for differential expression analysis of all RNA types, we used the known sample mix ratios and the fold-change measurements between samples A and B to construct the expected fold-changes between samples C and D for every gene [18]. By comparing the measured to the expected fold-changes between samples C and D, we found that both alignment-based pipelines showed superior performances over alignment-free pipelines (*R*^2^ = 0.62, 0.61, 0.46, and 0.42 for HISAT2+featureCounts, TGIRT-map, Kallisto, and Salmon, respectively; Fig. 4a). Further analyses showed that low correlations between measured and expected fold-changes were due to lowly-expressed genes (Fig. 4a), as suggested previously [15, 19]. However, to our surprise, Kallisto and Salmon had exceptionally poor ts to the corresponding models for these lowly-expressed genes (Fig. 4a; *R*^2^ = 0.44, 0.41, 0.13, and 0.06 for the lowest 75% expressed genes from HISAT2+featureCounts, TGIRT-map, Kallisto, and Salmon, respectively).

**Figure 4.**
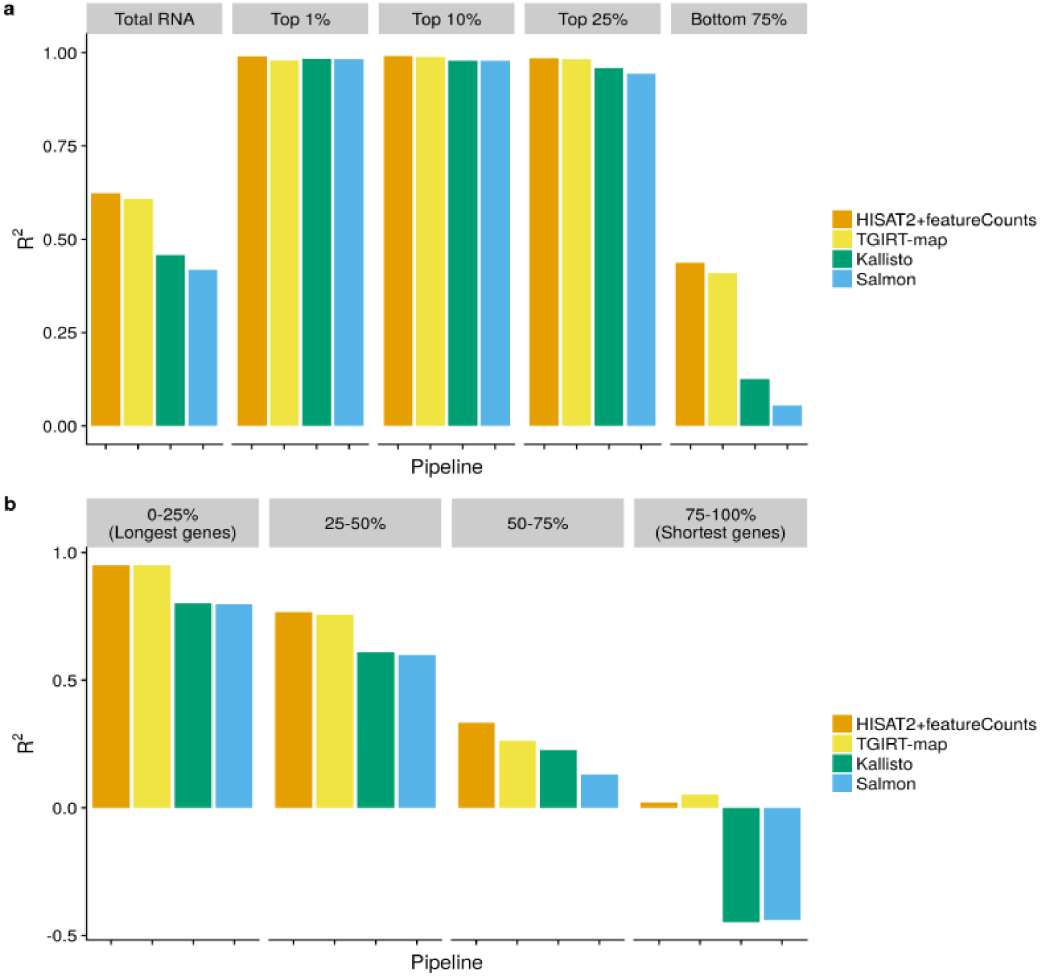
*R*^2^ of measured versus expected log2 fold-changes between samples C and D. (a) Gene expression level influenced accuracies of fold-change estimation. *R*^2^ values were computed from the expected and measured log2 fold-change values between samples C and D for each pipeline using different gene sets grouped by average gene expression levels. The first gene set, labeled “Total RNA”, includes all genes. The subsequent gene sets include only the genes with the top 1%, top 10%, top 25%, or bottom 75% expression levels, as indicated. Bars are color-coded by pipelines. (b) Gene lengths influenced accuracies of fold-change estimation. Genes from each pipeline were grouped by their gene lengths into four quantile groups. For each quantile group, *R*^2^ values were computed from the expected and measured log2 fold-change values between samples C and D. Bars are color-coded by pipelines. Coefficients of determination (R^2^) were computed by R2 function from *R* caret package [36]. Negative *R*^2^ values indicate exceptionally bad fold-change predictions [22] from the software as illustrated in Additional File 7, where the fold-change output do not fit well to the samples mix-ratio.

In addition to lowly-expressed genes, we also found that short gene lengths greatly decreased the accuracies in fold-change analyses, particularly for alignment-free pipelines (Fig. 4b; *R*^2^ = 0.02, 0.05, –0.45, and –0.44 for the shortest 25% genes from HISAT2+featureCounts, TGIRT-map, Kallisto, and Salmon, respectively). As we are testing whether the measured (observed) fold-changes fit into the expected fold-changes (model) constructed by the fold-changes between samples A and B and the known mix-ratios of samples A and B in samples C and D, *R*^2^ values served as a metric in quantifying the measurement errors relative to the known model. Thus, a negative *R*^2^ indicates highly discordant measurements relative to the expected fold-changes predicted by the known mix-ratios and the measured fold-changes between samples A and B [22], as illustrated in Additional File 7. These discordant measurements reect the incapabilities of the quantification tools in making accurate fold-change predictions either between samples A and B or between samples C and D for small genes.

To evaluate whether the deficiency in small genes quantification was specific to certain gene types, we computed the root mean square errors (RMSE) for the expected versus measured log2 fold-changes between samples C and D for genes grouped by their gene types. Consistent with the gene quantification results, Salmon and Kallisto showed slightly better fold-change estimation performances on ERCC spike-ins while alignment-based pipelines showed slightly better fold-change estimations on protein-coding genes (Fig. 5a and Table 1). Furthermore, alignment-based and alignment-free pipelines produced similar results for the majority of small RNA types, such as snoRNAs. For snoRNAs and other small non-coding RNAs (labeled as other sncRNA), Salmon and TGIRT-map recovered more genes than did HISAT2+featureCounts and Kallisto, but Salmon and TGIRT-map also showed similar RMSE values that were comparable to the RMSE values from HISAT2+featureCounts and Kallisto (Fig. 5a and Table 1). However, for tRNAs, alignment-based pipelines detected higher numbers of tRNA isoacceptors compared to alignment-free pipelines and showed advantages in detecting differential tRNA expression, yielding lower RMSE values (Fig 5b and Table 1).

**Figure 5.**
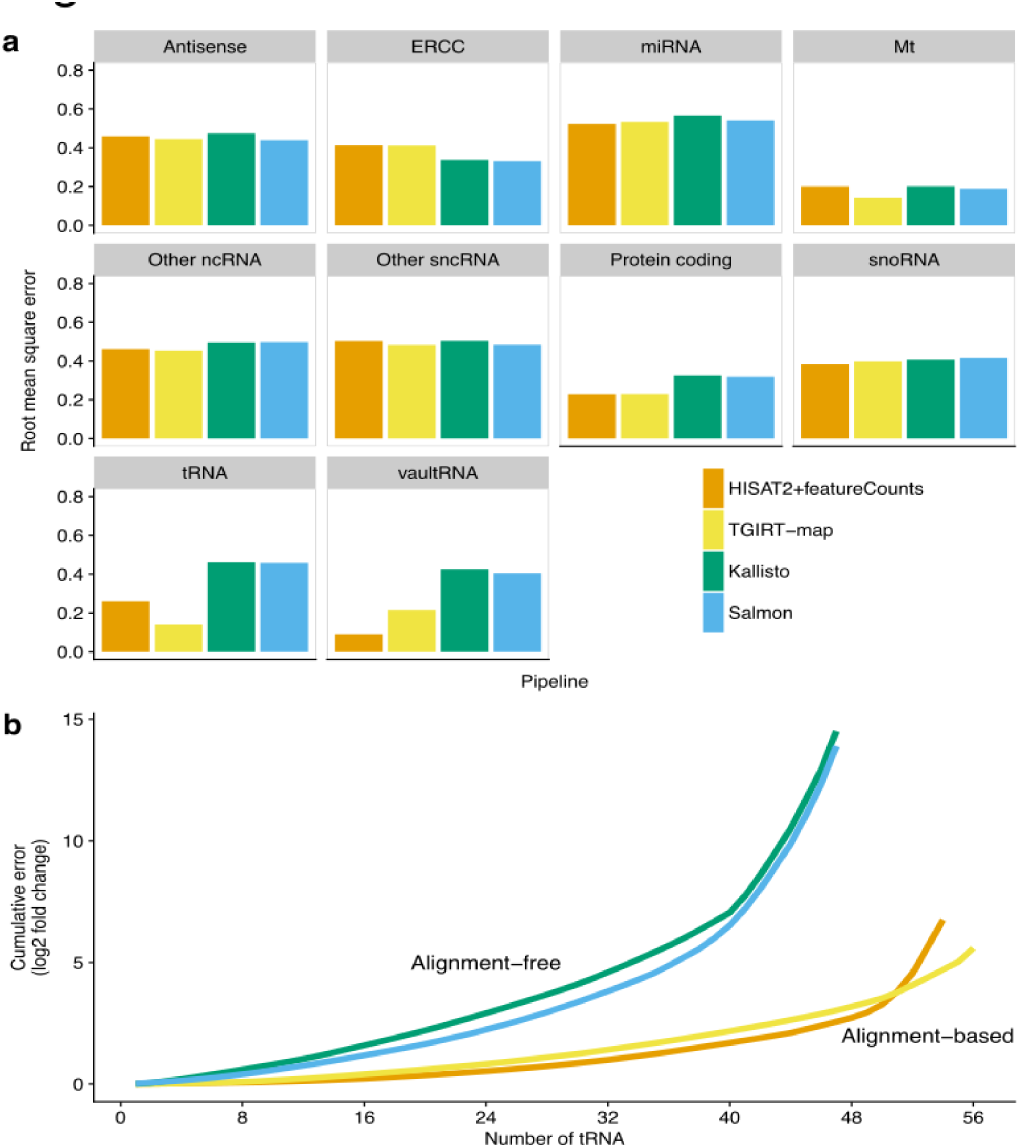
Pipeline performances of differential expression measurements for different gene types. (a) Root mean square error (RMSE) values between measured and expected log2 fold-change values for each gene type. Bars indicate RMSE values for different gene types from each pipeline and are color-coded by pipelines. (b) Cumulative absolute errors in tRNA log2 fold-change predictions. tRNAs (x-axis) were ordered ascendingly by the absolute errors in log2 fold-change predictions for each pipeline. Cumulative absolute errors for all detected tRNAs are plotted. Lines are color-coded by pipelines.

**Table 1.**
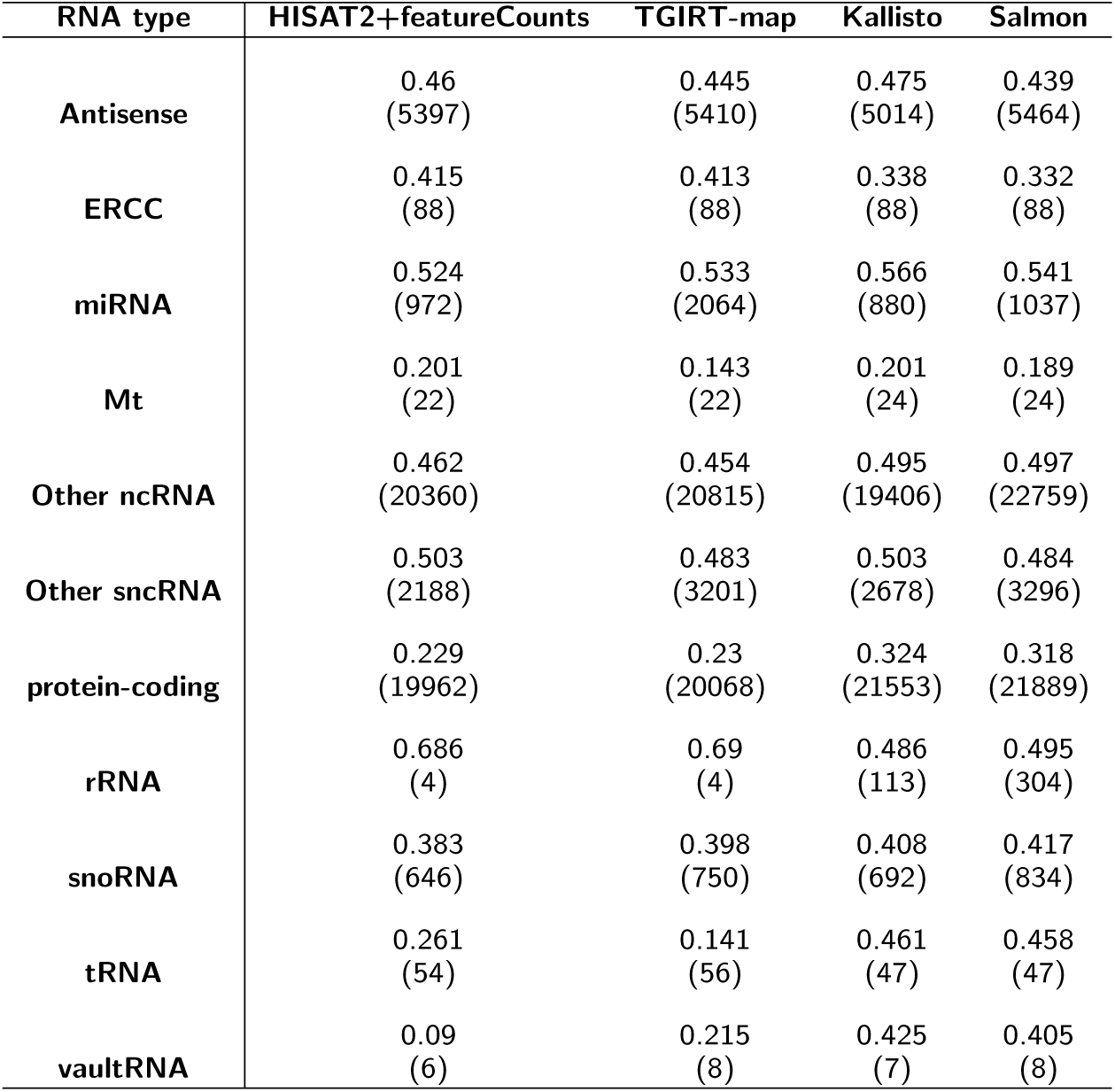
RMSE values in differential expression measurements. Numbers of detected genes are shown in parentheses. RNA-type annotations were generated from Ensembl [28]. “Other ncRNA” represents the following RNA types: sense_intronic, 3prime_overlapping_ncRNA, processed_transcript, sense_overlapping, lincRNA, and all pseudogenes. "Other sncRNA" represents the following RNA types: misc_RNA, snRNA, scaRNA, sRNA, scRNA. Mt represents all mitochondrial genes, including mitochondrial-encoded tRNAs. vaultRNA represents any Ensembl gene names with Vault or VTRNA.

As we showed distinct performances in tRNA fold-change analysis between alignment-free and alignment-based pipelines and a performance decrease in small gene quantification for alignment-free pipelines, we anticipated the comparison in tRNA fold-change predictions among the different pipelines would give us insights into the deficiency of alignment-free pipelines in small gene quantification. We hypothesized that the deficiency of alignment-free pipelines in small gene quantification was possibly due to the choice of a long k-mer size (31-mer) relative to the sizes of the small RNA transcripts, such as 75 nt for tRNAs. To investigate whether the choice of k-mer size had any effect on small RNA quantifications, we tested Salmon with four different k-mer sizes, ranging from 11 to 31 (11, 15, 21, and 31; default is 31 for both Salmon and Kallisto). We found that at *k* = 21, higher *R*^2^ values were observed for total RNA (Additional Files 8). Using tRNA as a model for comparisons in small gene quantifications, we also found a performance improvement (detected higher number of tRNA and lower RMSE) using *k* = 21. However, Salmon with this k-mer size still yielded two-fold higher RMSE relative to TGIRT-map for tRNA fold-change prediction (Additional Files 2 and 8).

To gain better understanding to the problem of small gene quantification in alignment-free pipelines, we further inspected the tRNAs that were undetected by both Salmon and Kallisto. Using TGIRT-map alignment results, we found that the read alignments mapped to these tRNAs displayed a high abundance of non-reference bases (Additional File 9). These non-reference bases may be caused by post-transcriptionally modified RNA bases that could introduce reverse-transcription errors by changing base-pairing interfaces [23, 24], as it is known that tRNAs are post-transcriptionally-modified and have abundant base modifications [15, 17, 25]. Thus, we predict that these abundant non-reference bases in small RNAs, tRNA in this case, may have prevented k-mer-based algorithms from successfully counting these reads.

## Discussion

We have performed an in-depth comparison among four RNA-seq pipelines, including two alignment-based and two alignment-free pipelines, to determine the relative performance of these tools for simultaneous quantification of long and small RNAs. The two alignment-based pipelines that we have tested were a widely-adopted align-and-count pipeline (HISAT2+featureCounts) [3, 9] and a custom gene-counting procedure with multi-step iterative alignments (TGIRT-map; Additional File 1); the two tested alignment-free pipelines were k-mer counting tools with and without bias corrections (Salmon [6] and Kallisto [5]). We have tested these four pipelines on quantification of both long and small RNAs in a novel benchmarking dataset generated by a thermostable group II intron reverse transcriptase (TGIRT) [15]. This dataset is unique in that small non-coding RNAs are highly represented, while the quality of long RNA quantification is comparable to that of other widely-adopted RNA-seq methods [15]. Using this dataset, we have found that while all four pipelines perform similarly on long and highly abundant RNAs, alignment-free tools have clear limitations for small and lowly-expressed RNA quantification.

For long gene quantification, we have found that all four pipelines quantify common gene targets (e.g. ERCC spike-ins and protein-coding genes) with similar results, confirming a previously benchmark study on poly-A selected RNA-seq [13]. Generally, gene quantification tends to be more similar between pipelines of the same type (i.e., HISAT2+featureCounts and TGIRT-map, or Kallisto and Salmon) than pipelines of the other type (alignment-based versus alignment-free). This result further supports a previous finding which showed more similar transcript isoforms quantifications were found among alignment-free or among alignment-based pipelines than among pipelines of the different types [14], suggesting that alignment-based pipelines may have somewhat different quantification biases than do the probabilistic models of alignment-free pipelines. Regardless of the different in gene quantifications, our results on differentially-expressed gene detections for long genes have supported that all four pipelines performed comparably for ERCC spike-ins and protein-coding genes when compared to their expected fold-changes (ERCC spike-ins) [21] or MAQC TaqMan assay measurements (protein-coding genes) [18]. This result further confirmed previous benchmark studies where alignment-free and alignment-based tools gave similar results differentially-expressed gene detection [13, 14].

For total gene quantification and differential expression analysis, all tested pipelines generally have performed similarly, with most disagreements occurring between pipeline comparisons of different pipeline types (i.e. an alignment-based pipeline vs an alignment-free pipeline). In the analyses of genes that were inconsistently quantified among pipelines, our results have confirmed that both high gene expression and long gene length were crucial to consistent abundance estimation, as suggested previously [13, 26]. Using fold-change analyses for comparisons in quantification accuracies, we have found that alignment-based tools were more accurate in quantifying lowly-expressed or small genes. This result likely reflects the nature of probabilistic assignments of k-mers and the inferences of TPM values. Although we have found that alignment-free pipelines were unreliable for quantifying extremely small RNAs (with shorter gene lengths) in total transcriptome analysis, alignment-free tools have performed comparably to alignment-based tools for most of the small RNA types, such as snoRNAs or other sncRNAs, in differential expression analyses. This disagreement between small non-coding RNAs and small mRNAs is possibly due to their differences in secondary structures and their different sensitivities to RNA fragmentations prior to RNA-seq library constructions, which over-fragmented mRNA fragments may be dropped-out from alignment-free quantifications.

Even though we have found that all pipelines perform similarly on the majority of small non-coding RNAs, our results have revealed that pipelines involving genome-alignment steps were superior to alignment-free tools in quantifying tRNAs speci-fically. We initially hypothesized the differences in performance were due to the choice of k-mer size in alignment-free pipelines and we have found an improvement in small gene quantifications when a moderately smaller *k* was chosen (*k* = 21). However, we have also found that the performance of alignment-free quantification at this optimal *k* was still not comparable to alignment-based pipelines for small genes. Using tRNA as a model, our results have suggested that this performance difference was likely due to a combinatorial effect of the choice of k-mer size and misincorporations introduced by post-transcriptionally-modified RNA bases during reverse transcriptions. We reason that a relatively long k-mer and erroneous sequencing reads likely impede matching to the indexed transcriptome even when these sequencing reads are shredded into k-mers since all k-mers inherit the same errors/mismatches. Since mismatches on sequencing reads can either be reverse transcription errors or biological variations on small RNAs [27], we expect the same phenomenon may occur if the small RNAs contain single-nucleotide polymorphisms or mutations.

## Conclusions

In summary, we have shown that different types of pipelines performed similarly for common differentially-expressed gene targets such as protein-coding genes. However, accurate quantification of lowly-expressed or small RNA is difficult to achieve with alignment-free pipelines. Using tRNA as a model, we have also found that k-mer counting algorithms are not compatible for quantifying small genes with abundant mutations regardless of the choice of k-mer size.

## Abbreviations

TGIRT: Thermostable group II intron reverse transcriptase
MAQC: microarray quality control consortium
ERCC: External RNA controls consortium
RMSE: root mean square error
ROC: receiver operating characteristic
AUC: area under the curve
TPM: transcripts per million

## Methods

### Data and reference preparation

Raw sequencing reads for TGIRT-seq data generated from the well-studied MAQC samples were downloaded from the NCBI Sequence Read Archive, accession number SRP066009 [15]. In brief, this dataset includes triplicates of four different human total RNA samples (called A{D) spiked with External RNA Control Consortium (ERCC) transcripts. ERCC spike-in transcripts are 92 *in* vitro transcripts with 250-2000-nt long [21]. Two ERCC spike-in mixes (mixes 1 and 2) with different concentrations for each transcript were spiked into RNA samples A (universal human RNA) and B (human total brain RNA), respectively, to provide known fold-changes of these spike-in transcripts between these two samples [18, 21]. By design, these two spike-in mixes establish four different differentially-expressed gene groups with relative ratios of 0.67:1, 1:1, 1:2, and 1:4 between samples A and B. Samples C and D are mixes of samples A and B in ratios of 3:1 and 1:3, respectively. For detailed library preparations, please refer to Nottingham, *et al* [15].

Human transcriptome reference sequences were obtained from Ensembl (human genome build version GRCh38.87) by downloading cDNA and non-coding RNA sequence FASTA les [28]. ERCC spike-in sequences were downloaded from the vendor’s website (ThermoFisher). Transfer RNA (tRNA) annotations and sequences were downloaded from the Genomic tRNA Database [29]. For tRNA sequences, introns were removed, and all sequences were de-duplicated at the sequence level to reduce multiply-mapped reads and improve tRNA counting. 5S rDNA (GenBank accession: X12811.1) and complete rDNA repeat unit (GenBank accession: U13369.1) sequences were downloaded from GenBank (NCBI). These sequences were merged and indexed with Kallisto v0.43.0 [5] and Salmon v0.8.2 [6] using the default k-mer size (*k* = 31). Human genome sequences and annotations were downloaded as FASTA and GTF files from Ensembl (human genome build version GRCh38.88) [28], combined with ERCC spike-in sequences and rDNA sequences, and indexed by HISAT2 v2.0.4 [9]. A HISAT2 splice-site file was created using the human genome build GRCh38.88 GTF file [9]. Gene annotations were obtained as a BED file using Ensembl biomaRt package in R [30, 31] from Genome build version GRCh38.87 [28]. The gene annotation BED file was merged with tRNA, ERCC spike-in, and rDNA annotations for gene counting. The merged gene BED file was converted to a featureCounts SAF file [3] using the UNIX awk command.

### Adaptor trimmings

Raw reads were adapter- and quality-trimmed by cutadapt v1.13 [32] via:

~~~
cutadapt -m 15 -O 5 -n 3 -q 20 \
     -b AAGATCGGAAGAGCACACGTCTGAACTCCAGTCAC \
     -B GATCGTCGGACTGTAGAACTCTGAACGTGTAGA \
     -o ${TRIMMED_FQ1}-p ${TRIMMED_FQ2}\
     ${FQ1} ${FQ2}
~~~

### HISAT2 mapping

Reads were aligned to the genome using HISAT2 v2.0.4 [9] via:

~~~
hisat2 -k 10 ‐‐no-mixed -no-discordant \
   ‐‐known-splicesite-infile={SPLICESITE_FILE} \
   -1 ${TRIMMED_FQ1} -2 ${TRIMMED_FQ2}
~~~

### HISAT2+featureCounts

Trimmed reads were aligned using HISAT2 mapping. Gene counts from HISAT2 mapping were generated by featureCounts v1.5.0 [3] via:

~~~
featureCounts -F SAF -O -s 1 -M -T 24 \
       ‐‐largestOverlap ‐‐minOverlap 10 \
       ‐‐primary -p -P -d 10 -D 10000 -B \
       -S fr -C -donotsort \
       ${LIST_OF_BAM_FILES}
~~~

### TGIRT-map

Our custom pipleine TGIRT-map first filters out all trimmed reads that can be aligned to tRNA or rRNA references to reduce multiply-mapped reads (Additional File 1). The remaining unaligned reads are then sequentially aligned to the human genome using a splice-aware aligner (HISAT2) [9] and a sensitive local aligner (BOWTIE2) [20]. A single alignment locus is picked from the multiply-mapped fragments by asserting the assumptions that (A) RNA-seq fragments are small (smallest insert size), (B) ribosomal genes are abundant in the genome (ribosomal gene loci), and (C) fragments are unlikely to be originated from haplotype or patch sequences (as defined by Ensembl annotations) of the artificially assembled genome. Finally, gene quantification is done on genomic loci, except for tRNA and rRNA which require an additional step of re-aligning and counting.

Trimmed reads were first aligned to rRNA (GenBank accession numbers: X12811.1 and U13369.1) and tRNA sequences with BOWTIE2 v2.3.2 [20] via:

~~~
bowtie2 -D 20 -R 3 -N 0 \
        -L 8 -i S,1,0.50 ‐‐no-mixed \
        ‐‐norc ‐‐no-discordant \
        -1 ${TRIMMED_FQ1} -2 ${TRIMMED_FQ2}
~~~

We define the mapped reads as tRNA and rRNA pre-mapped reads. Unmapped reads were then aligned to the genome with HISAT2 as described above in HISAT2 mapping [9]. Unaligned reads from HISAT2 mapping were then rescued by realigning locally to the genome with BOWTIE2 [20] via:

~~~
bowtie2 -D 20 -R 3 -N 0 -L 8 \
     -i S,1,0.50 -p 24 -k 10 \
     ‐‐no-mixed ‐‐no-discordant \
     -1 ${FQ1} -2 ${FQ2}
~~~

All alignment pairs with > 10 nucleotides of soft-clipped bases on either reads or were discordant pairs were discarded. Multiply-mapped reads from the two genome-mapping steps were grouped and a pair of best alignment was chosen by the following ordered criteria: the alignment pair (A) had the smallest insert size, (B) was mapped to ribosomal gene locus, or (C) was mapped to chromosome 1-23, X or Y. If none of these criteria was met by a single alignment pair, a random alignment pair was chosen from the multiply-mapped loci. These filtered alignments were then merged with the uniquely-mapped alignments. All reads that mapped to tRNA or rRNA loci were extracted by BEDtools v2.26 using pairtobed command with options -s -f 0.01 [33], combined with tRNA and rRNA pre-mapped reads and re-aligned to tRNA and rRNA references. Counts were generated from the aligned BAM file. Counts for each anticodon were aggregated. Other gene counts were calculated by converting the genome alignments to fragment coordinates in a BED file using the BEDtools bamtobed command and counted using the BEDtools coverage command [33]. TGIRT-map pipeline is available at: https://github.com/wckdouglas/tgirt_map.

### Kallisto and Salmon

We used Kallisto v0.43.0 [5] and Salmon v0.8.2 [6] for our alignment-free pipelines. In both cases, adaptor-trimmed reads were used as input.

We called Salmon with the following command-line arguments:

~~~
salmon quant \
       ‐‐seqBias ‐‐gcBias \
       ‐‐index ${INDEX} ‐‐libType ISF \
       ‐‐writeMappings \
       ‐‐threads=${THREADS} \
       ‐‐auxDir ${RESULTPATH} \
       ‐‐numBootstraps 100 \
       ‐‐mates1 ${TRIMMED_FQ1} ‐‐mates2 ${TRIMMED_FQ2} \
       ‐‐output ${RESULTPATH}
~~~

We called Kallisto with the following command-line arguments:

~~~
kallisto quant \
      -i ${INDEX} -o ${OUTPUT_PATH} \
      ‐‐fr-stranded ‐‐threads=${THREADS}\
      ${TRIMMED_FQ1} ${TRIMMED_FQ2}
~~~

### Differential expression analysis

DESeq2 v1.14.1 [34] was used for all differential expression analyses.

Because DESeq2 does not accept TPM values as input, transcript TPM values from Salmon and Kallisto were converted to gene-level counts using Tximport [35] before any differential-expression analyses.

## Availability of data and material

All scripts used for data processing are deposited on GitHub at: https://github.com/wckdouglas/tgirt_benchmark.

## Author’s contributions

D.C.W., A.M.L., C.O.W. designed the experiments. D.C.W., K.S.H., J.Y., A.M.L. and C.O.W. wrote the manuscript. D.C.W., K.S.H and J.Y. conducted the experiments and analyzed the data with input from A.M.L. and C.O.W.

## Acknowledgements

We thank Dr. Nicholas C. Wu (The Scripps Research Institute) for critical reading of the manuscript, and Ben R. Jack and Dr. Adam J. Hockenberry from the Wilke lab for fruitful discussions and useful insights. The Texas Advanced Computing Center (TACC) and the Center for Biomedical Research Support (CBRS) at the University of Texas at Austin provided high-performance computing resources.

## Additional Files

**Additional File 1.**
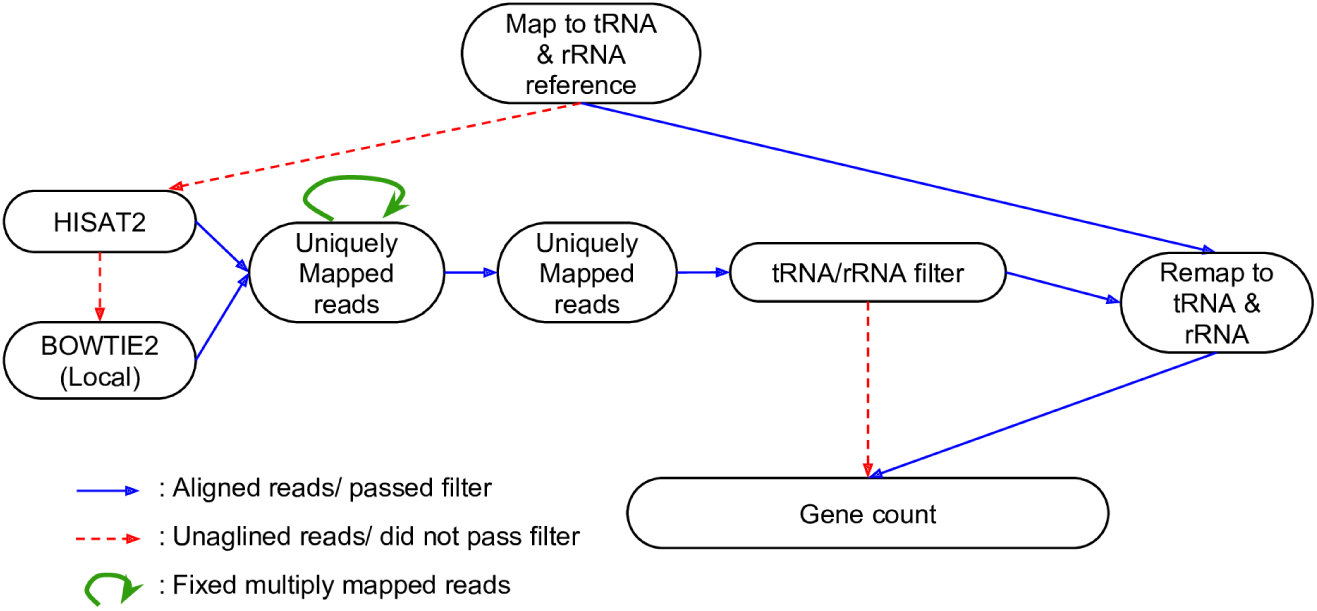
Flow chart of the custom pipeline TGIRT-map. As TGIRT-seq simultaneously recovers a large fraction of small structural RNAs in addition to regular long RNAs [15, 17], TGIRT-map was designed to optimize the quantification of these small genes. The pipeline is similar to the Genobee-exceRpt small RNA-seq pipeline [37], where reads are first aligned against the tRNA and rRNA sequences to avoid ambiguous assignments in later steps. Unaligned reads (red arrow) are iteratively aligned to the human genome by HISAT2 [9] and BOWTIE2 [20] to minimize unassigned reads. Multiply-mapped reads are extracted and assigned as uniquely-mapped at loci selected by the following criteria in order: the pair of alignments (A) has the smallest insert size, (B) is mapped to ribosomal gene loci, and (c) is mapped to chromosome 1-23, X or Y, otherwise a random pair of alignments is selected. From these genome alignments, reads that aligned to tRNA or rRNA loci are extracted and combined with tRNA and rRNA reads from the first step. Finally, these combined reads are realigned to the tRNA and rRNA sequences to generate gene counts.

Additional File 2—supplementary tables.zip Zip file containing several data tables in tab-separated format, as well as a readme file that explains the contents of each data file.

**Additional File 3.**
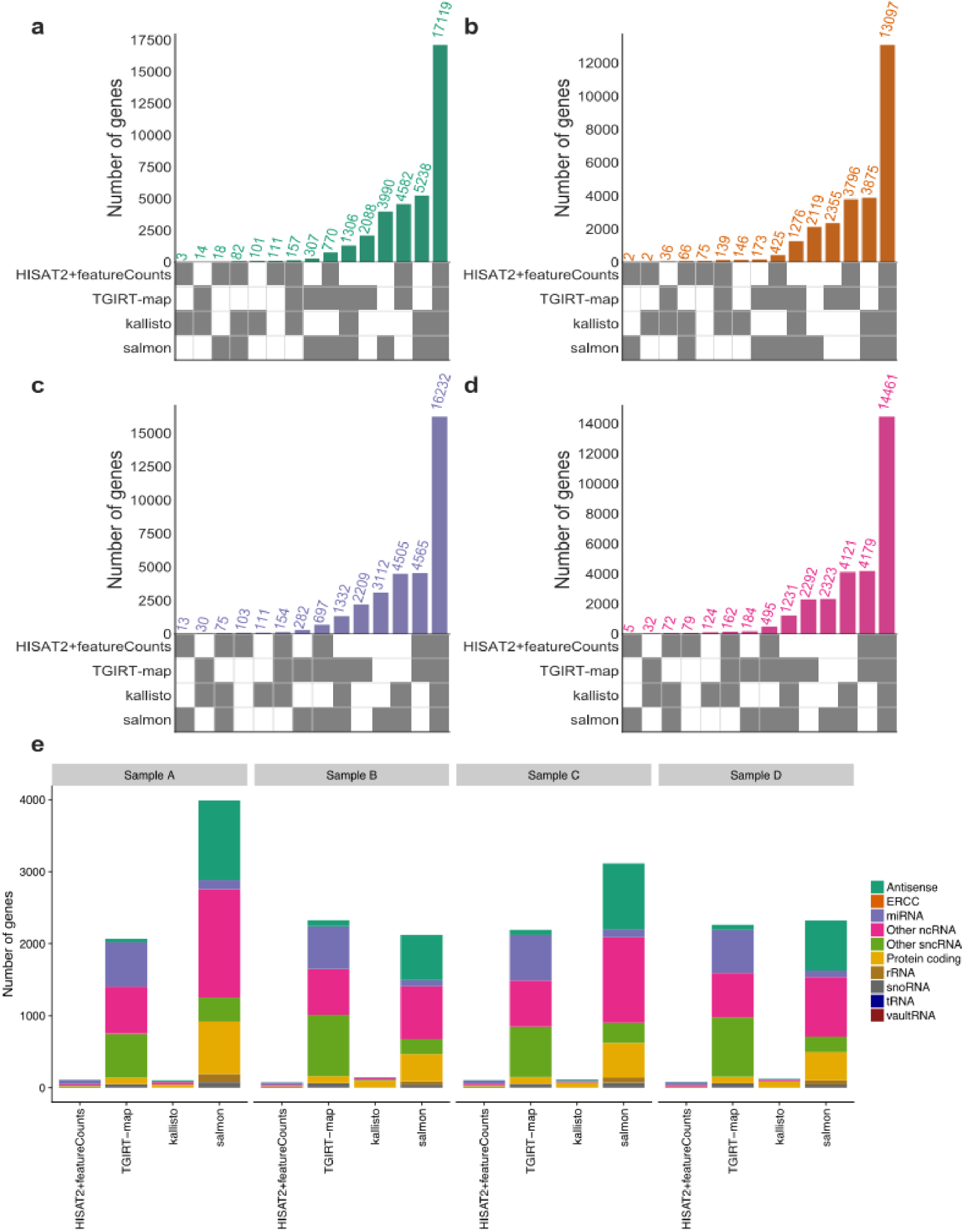
Intersections of detected genes among tested pipelines. Genes with TPM > 0.1 were defined as being detected. (a-d) UpSet graphs of intersections of detected genes among pipelines. An UpSet graph describes intersections of detected genes among subsets of tested pipelines. Each UpSet graph contains a bar chart and a sample-comparison matrix. Bars indicate the numbers of detected genes in the specific intersection specified in the sample-comparison matrix. Each bar indicates the number of genes detected only in the intersection of the set of compared pipelines. The sample-comparison matrix at the bottom describes the set of pipelines within which the comparison is made. Grey boxes in the matrix annotate the pipelines included in the comparison. (e) Uniquely-detected genes. The heights of the stacked bars indicate the numbers of genes uniquely detected by the corresponding pipeline. Stacked bars are color-coded with gene types.

**Additional File 4.**
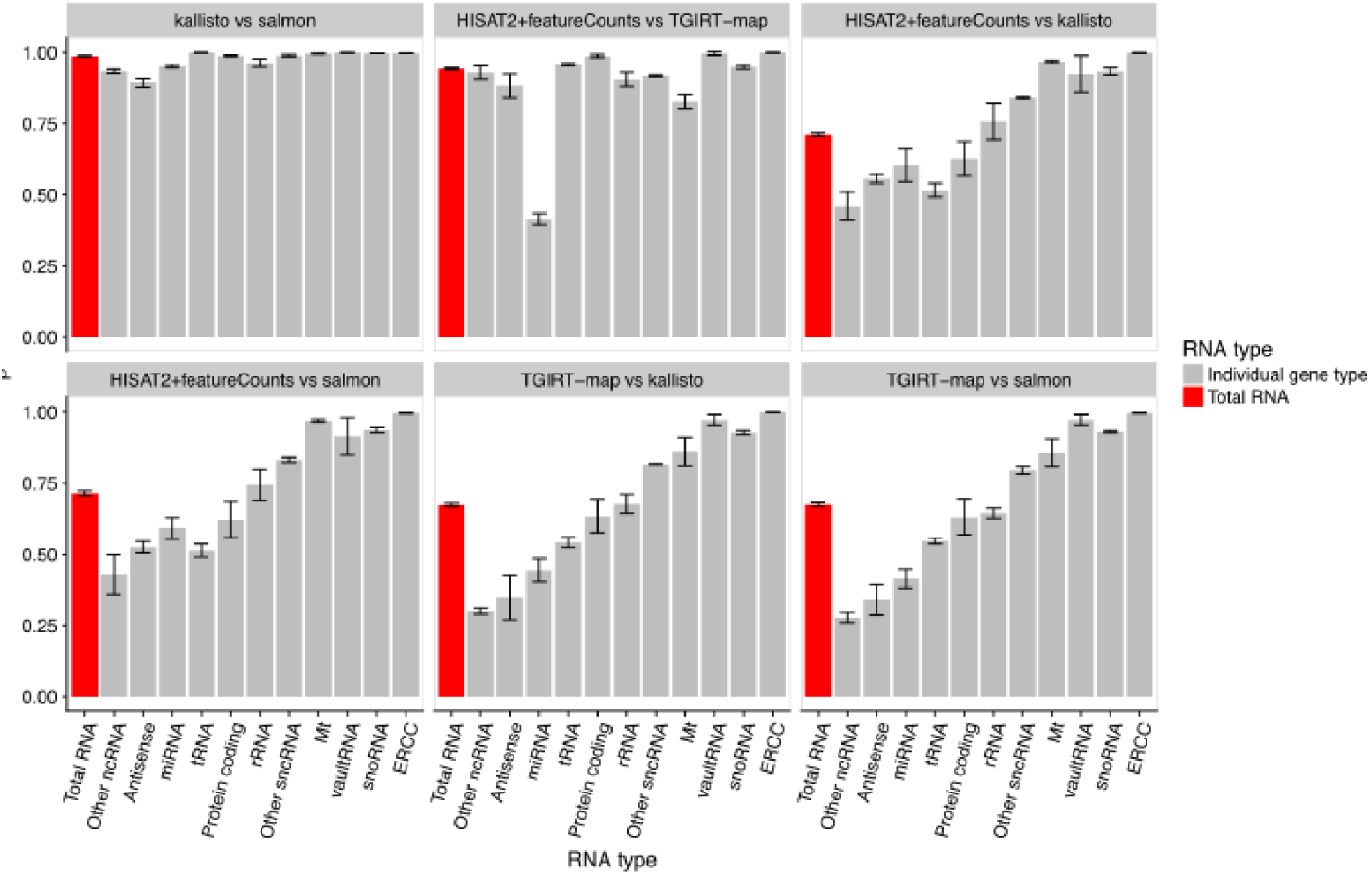
Pearson’s correlation coefficients for pairwise pipeline comparisons in gene quantifications. For every pairwise pipeline comparison and each sample (A{D), Pearson’s correlation coefficients were computed using the average log2 TPM of each gene across the triplicates for each sample. Bar heights indicate average correlations among the four samples A-D. Error bars represent standard deviations among the four samples A-D. Red bars represent total RNA correlation coefficients (all genes). Grey bars indicate correlation coefficients grouped by gene type. Each panel represents a single pairwise pipeline comparison.

**Additional File 5.**
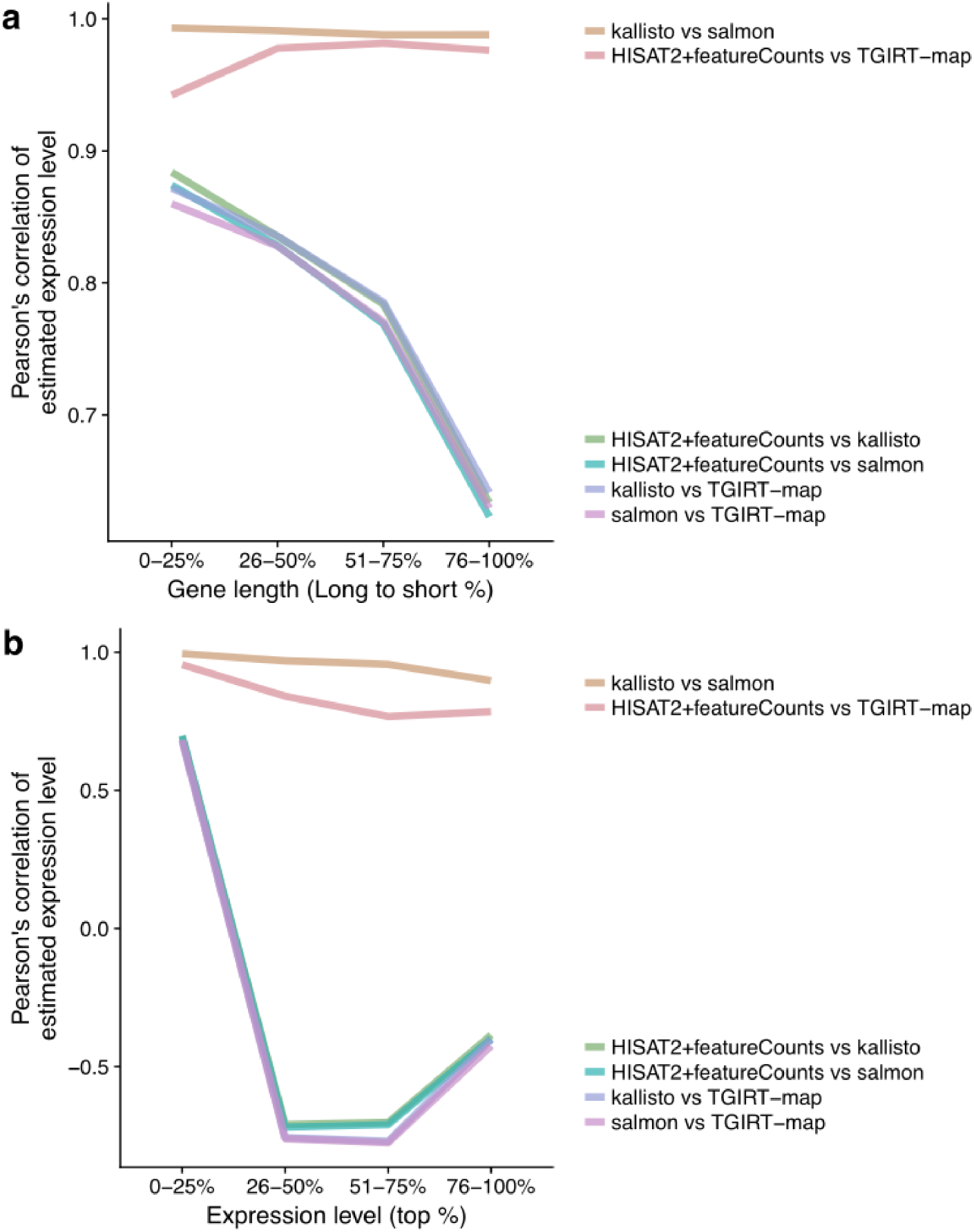
Impact of gene lengths and expression levels on gene quantification correlations among pipelines. (a) Gene lengths inuence gene expression estimation correlations. Genes were separated into quantile groups according to their lengths. For every quantile group from each sample, Pearson’s correlation coefficients of average TPMs for each gene across replicates were computed for every pairwise pipeline comparison and then averaged across the four samples. Plotted are the average correlations versus the quantile groups. Each line represents a pairwise comparison and is color-coded accordingly. (b) Gene expression levels influence gene expression estimate correlations. Genes were separated into quantile groups according to their average expression values (TPMs). For every quantile group from each sample, Pearson’s correlations were computed in the same way as in panel (a). Each line represents a pairwise comparison and is color-coded accordingly.

**Additional File 6.**
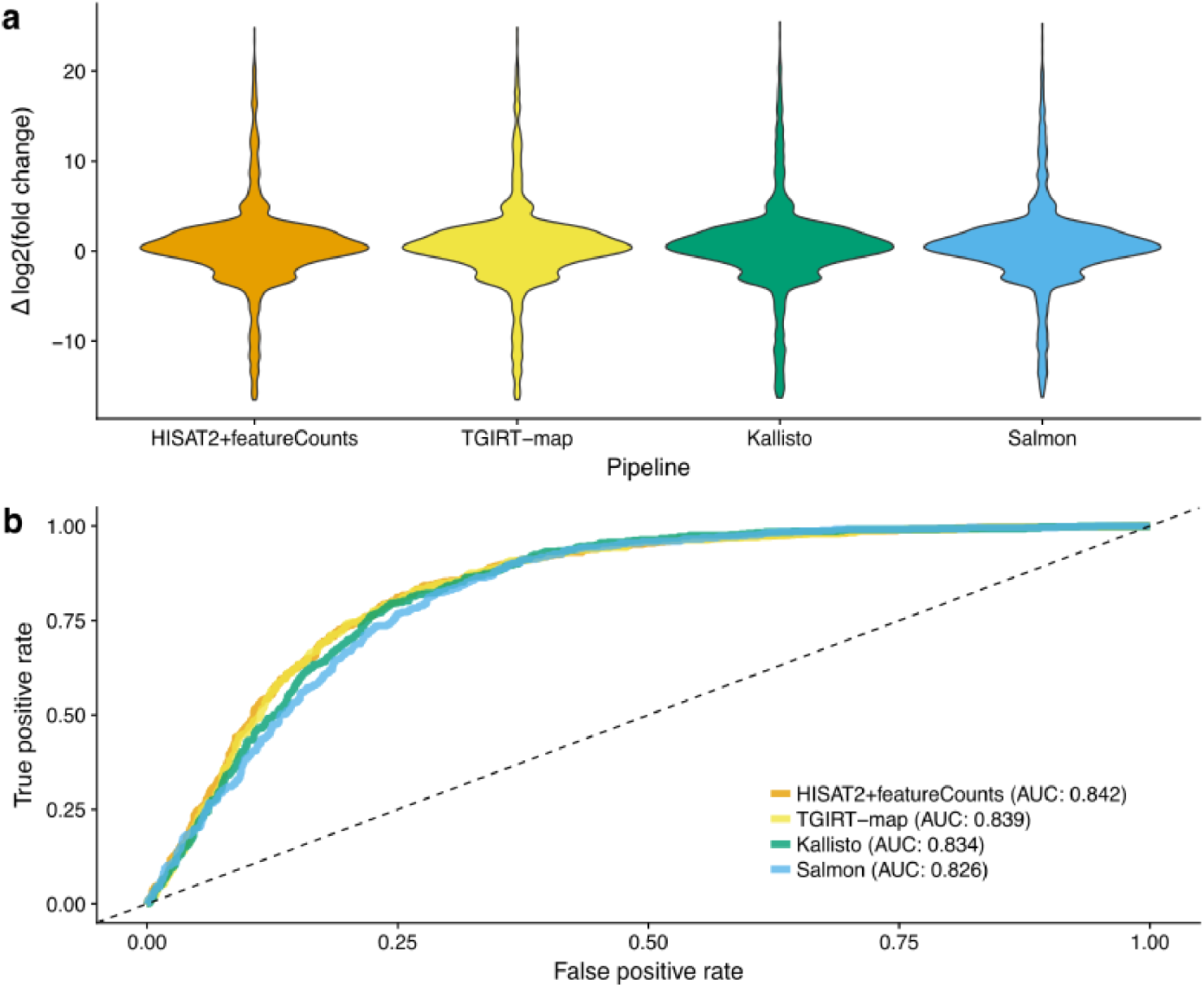
Differential expression analysis of protein-coding genes. (a) Comparison of measured fold-changes between samples A and B from the TGIRT-seq analysis and the MAQC TaqMan assay. MAQC TaqMan assay data for the same samples were downloaded from the NCBI Gene Expression Omnibus (GEO), accession number GSE5350 [18]. 972 unique protein-coding genes were compared. Log2 fold-changes were calculated from MAQC data and compared to the DESeq2 results for each pipeline. Distributions of the deviations between RNA-seq and MAQC data log2 fold-changes were plotted as violin-plots. Distributions are color-coded by pipelines. (b) ROC curves for calling differentially-expressed (DE) protein-coding genes. MAQC TaqMan assay genes were binary-labeled as DE if absolute log2 fold-change >0.5 and not DE otherwise (739 DE and 225 non-DE), as suggested by a previous study [10]. ROC curves for DESeq2 results from abundance estimates computed by each pipeline were plotted. AUCs were computed for each ROC curve to quantify performances of the differential expression caller (DESeq2) [34] on abundance estimates from each pipeline. ROC curves are color-coded by pipelines. The dotted diagonal line indicates random guessing (AUC=0.5).

**Additional File 7.**
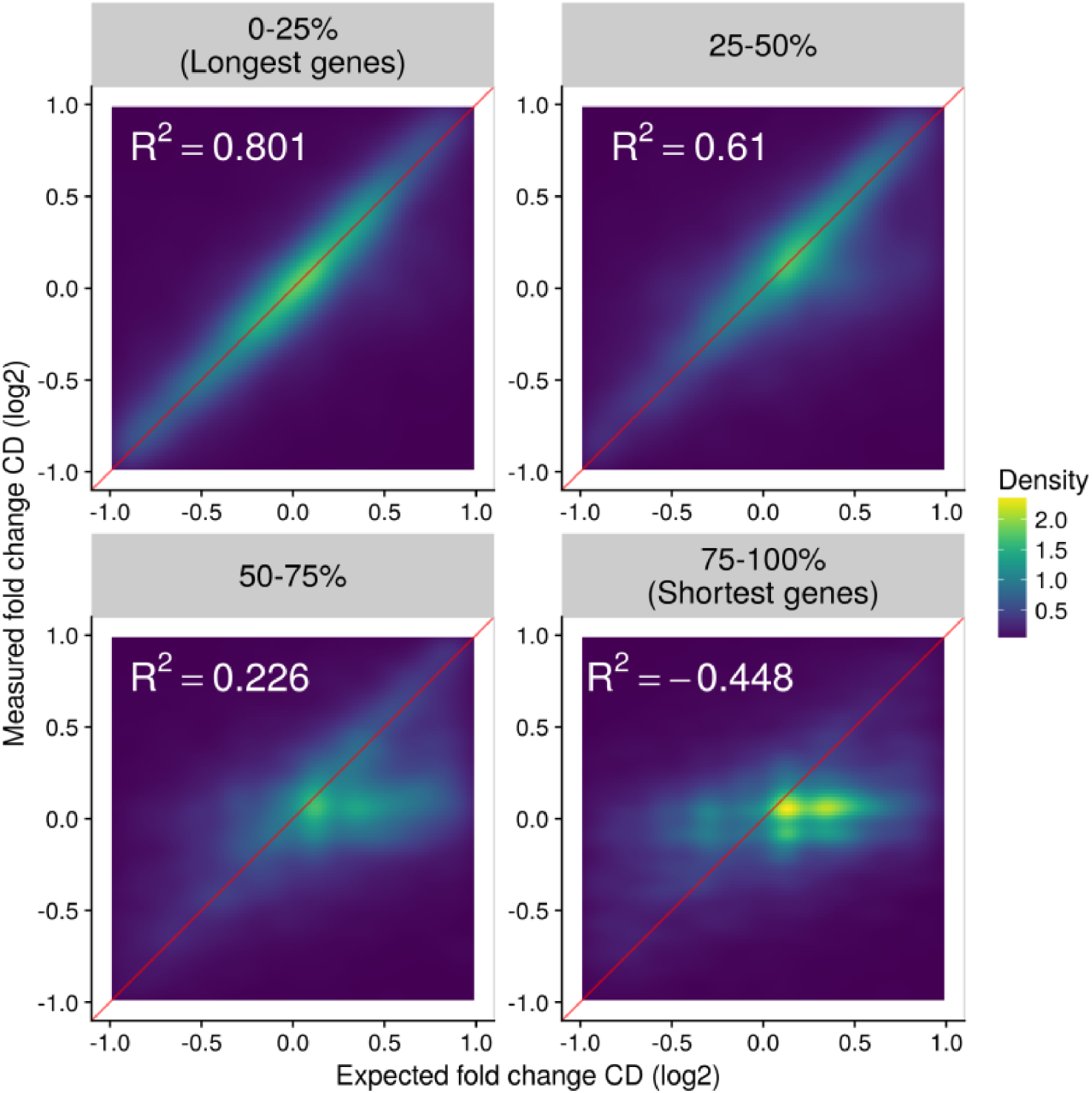
Measured versus expected log2 fold-changes between samples C and D for Kallisto. Expected log2 fold-changes for each gene (denoted by red line) were constructed from the (A) Kallisto-measured log2 fold-changes between samples A and B and (B) the known sample mix ratio of samples A and B in samples C and D, using a previously-described algorithm [18]. Each panel shows a two-dimensional kernel density estimation of the distribution of measured vs expected log2 fold-changes between samples C and D for genes grouped by quantile groups of gene lengths. Coefficients of determination for each group of genes are annotated as *R*^2^. Perfectly correlated log2 fold-changes are annotated by the diagonal red lines. Panels with high *R*^2^ generally show denser trends along the red line, indicating the log2 fold-changes between samples C and D were accurately predicted by the software that recapitulated the sample-mixing ratio and thus, fitted better to the model (red line). For groups with lower *R*^2^, a denser (lighter color) horizontal trend was observed on the right side of the red line, indicating the software either over-estimated log2 fold-changes between samples A and B that filed to construction of an incorrect model or under-estimated log2 fold-changes between samples C and D that filed to a poor fit to the model.

**Additional File 8.**
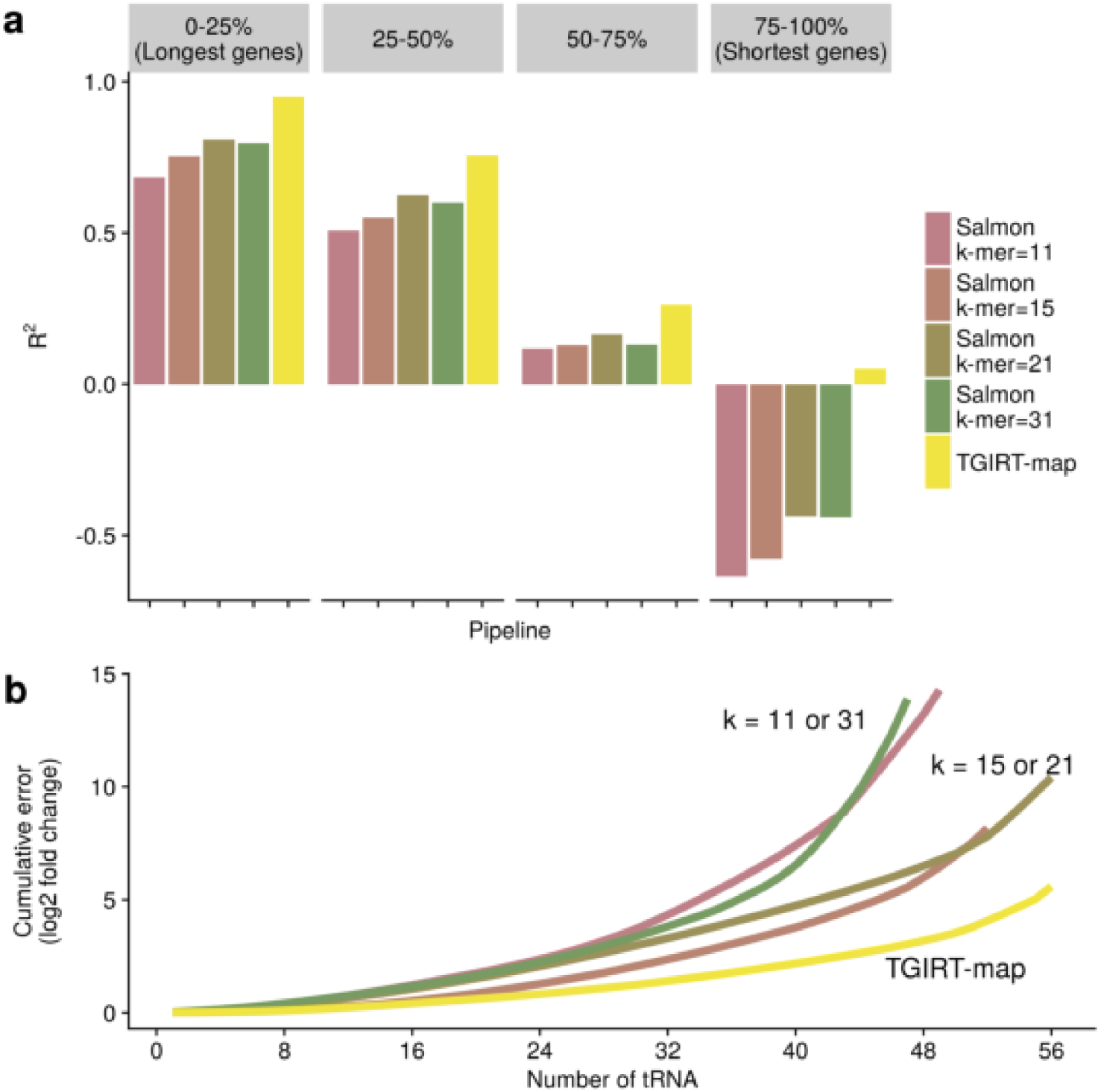
Performances of Salmon using different k-mer sizes of 11, 15, 21, and 31 (default). (a) Accuracies of fold-change estimation in quantile groups of gene lengths. All detected genes from TGIRT-map or Salmon with different k-mer sizes were grouped by their gene lengths into four quantile groups. For each quantile group, *R*^2^ values were computed from the expected and measured log2 fold-change values between samples C and D. Bars are color-coded by pipelines. Coefficients of determination (*R*^2^) were computed by R2 function from R caret package [36]. (b) Cumulative absolute errors in tRNA log2 fold-change predictions for each k-mer size and TGIRT-map. tRNAs were sorted in ascending order by their absolute errors in log2 fold-change predictions. Cumulative absolute errors for all detected tRNA were plotted. Lines are color-coded by pipeline.

**Additional File 9.**
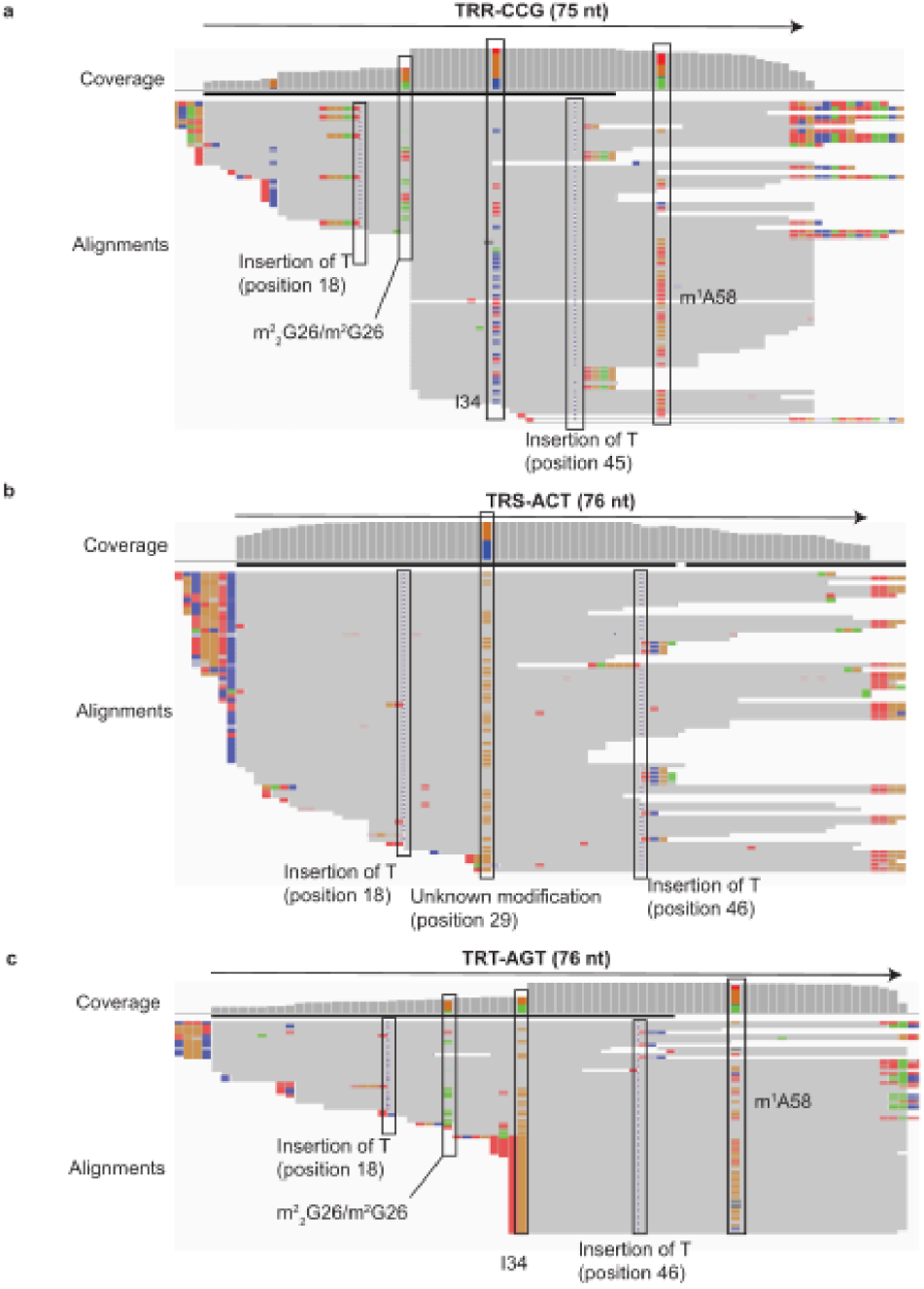
Erroneous sequencing reads may have impeded k-mer counting for tRNAs. IGV genome browser screen-shots of three tRNA alignments from Sample A1 aligned via TGIRT-map were shown. The three tRNA alignments being shown are (a) TRR-CCG, (b) TRS-ACT, and (c) TRT-AGT. Arrows on the top of each panel indicate the corresponding tRNA transcript strands (5’ to 3’) and sizes. Bar charts below the arrows represent coverages at every position along the tRNA transcripts. Read alignments from Sample A1 aligned by TGIRT-map are shown below the bar charts. Grey colors represent read bases that match the reference bases. Other colors indicate bases on the read alignments that do not match the reference bases (purple, red, green, blue, and brown colors indicate insertions, thymidines, adenosines, cytidines, and guanosines on the read alignments, respectively). These errors may reflect post-transcriptionally-modified RNA bases, which interfere with canonical base-pairings and filead to misincorporations during TGIRT reverse transcriptions [15, 17, 25]. Errors due to misincorporations at known and unknown sites of post-transcriptionally modified RNA bases are highlighted with boxes (m^2^ _2_G26: N2,N2-dimethylguanosine at position 26; m^2^G26: N2-methylguanosine at position 26; I34: Inosine at position 34; m^1^A58: 1-methyladenosine at position 58; see also [15, 17, 38]). Unannotated mismatches from both ends can be untrimmed adapters or tRNA-precusor sequences. We observe that on these tRNAs, the errors induced by RNA-base-modification occurred at positions ∼20, 30, or 50 on ∼75 nucleotide long tRNA transcripts. Therefore, we expect that every k-mer originating from an erroneous sequencing read inherits at least one of these errors, and therefore k-mers cannot be matched accurately to the transcript database. The effect is worse when a large k-mer size is selected, such as the default *k* = 31.

## References

1. Mortazavi, A., Williams, B.A., McCue, K., Schaeffer, L., Wold, B.: Mapping and quantifying mammalian transcriptomes by RNA-Seq. Nature Method 5(7), 621–628 (2008).

2. Trapnell, C., Roberts, A., Goff, L., Pertea, G., Kim, D., Kelley, D.R., Pimentel, H., Salzberg, S.L., Rinn, J.L., Pachter, L.: differential gene and transcript expression analysis of RNA-seq experiments with TopHat and Cufflinks. Nature Protocols 7 (3), 562–578 (2012).

3. Liao, Y., Smyth, G.K., Shi, W.: featureCounts: an efficient general purpose program for assigning sequence reads to genomic features. Bioinformatics (Oxford, England) 30 (7), 923–930 (2014).

4. Patro, R., Mount, S.M., Kingsford, C.: Sailfish enables alignment-free isoform quantification from RNA-seq reads using lightweight algorithms. Nature Biotechnology 32 (5), 462–464 (2014)

5. Bray, N.L., Pimentel, H., Melsted, P., Pachter, L.: Near-optimal probabilistic RNA-seq quantification. Nature Biotechnology 34 (5), 525–527 (2016)

6. Patro, R., Duggal, G., Love, M.I., Irizarry, R.A., Kingsford, C.: Salmon provides fast and bias-aware quantification of transcript expression. Nat Meth (2017)

7. Pertea, M., Kim, D., Pertea, G.M., Leek, J.T., Salzberg, S.L.: Transcript-level expression analysis of RNA-seq experiments with HISAT, StringTie and Ballgown. Nature Protocols 11 (9), 1650–1667 (2016)

8. Dobin, A., Davis, C.A., Schlesinger, F., Drenkow, J., Zaleski, C., Jha, S., Batut, P., Chaisson, M., Gingeras, T.R.: STAR: ultrafast universal RNA-seq aligner. Bioinformatics 29 (1), 15–21 (2013).

9. Kim, D., Langmead, B., Salzberg, S.L.: HISAT: a fast spliced aligner with low memory requirements. Nature Methods 12 (4), 357–360 (2015).

10. Rapaport, F., Khanin, R., Liang, Y., Pirun, M., Krek, A., Zumbo, P., Mason, C.E., Socci, N.D., Betel, D.: Comprehensive evaluation of differential gene expression analysis methods for. Genome biology 14(9), 95 (2013).

11. Teng, M., Love, M.I., Davis, C.A., Djebali, S., Dobin, A., Graveley, B.R., Li, S., Mason, C.E., Olson, S., Pervouchine, D., Sloan, C.A., Wei, X., Zhan, L., Irizarry, R.A.: A benchmark for RNA-seq quantification pipelines. Genome Biology 17(1), 74 (2016).

12. Baruzzo, G., Hayer, K.E., Kim, E.J., Di Camillo, B., FitzGerald, G.A., Grant, G.R.: Simulation-based comprehensive benchmarking of RNA-seq aligners. Nature Method 14(2), 135–139 (2017)

13. Everaert, C., Luypaert, M., Maag, J.L.V., Cheng, Q.X., Dinger, M.E., Hellemans, J., Mestdagh, P.: Benchmarking of RNA-sequencing analysis workflows using whole-transcriptome RT-qPCR expression data. Scientific Reports 7(1), 1559 (2017).

14. Sahraeian, S.M.E., Mohiyuddin, M., Sebra, R., Tilgner, H., Afshar, P.T., Au, K.F., Asadi, N.B., Gerstein, M.B., Wong, W.H., Snyder, M.P., Schadt, E., Lam, H.Y.K.: Gaining comprehensive biological insight into the transcriptome by performing a broad-spectrum RNA-seq analysis. Nature Communications 8(1), 59 (2017).

15. Nottingham, R.M., Wu, D.C., Qin, Y., Yao, J., Hunicke-Smith, S., Lambowitz, A.M.: RNA-seq of human reference RNA samples using a thermostable group II intron reverse transcriptase. RNA 22(4), 597–613 (2016)

16. Mohr, S., Ghanem, E., Smith, W., Sheeter, D., Qin, Y., King, O., Polioudakis, D., Iyer, V.R., Hunicke-Smith, S., Swamy, S., Kuersten, S., Lambowitz, A.M.: Thermostable group II intron reverse transcriptase fusion proteins and their use in cDNA synthesis and next-generation RNA sequencing. RNA 19(7), 958–70 (2013).

17. Qin, Y., Yao, J., Wu, D.C., Nottingham, R.M., Mohr, S., Hunicke-Smith, S., Lambowitz, A.M.: High-throughput sequencing of human plasma RNA by using thermostable group II intron reverse transcriptases. RNA 22(1), 111–128 (2016)

18. Consortium, M.: The MicroArray Quality Control (MAQC)-II study of common practices for the development and validation of microarray-based predictive models. Nature Biotechnology 28(8), 827–838 (2010).

19. SEQC/MAQC-III Consortium: A comprehensive assessment of RNA-seq accuracy, reproducibility and information content by the Sequencing Quality Control Consortium. Nature Biotechnology 32(9), 903–914 (2014)

20. Langmead, B., Salzberg, S.L.: Fast gapped-read alignment with Bowtie 2. Nature Method 9(4), 357–359 (2012)

21. Jiang, L., Schlesinger, F., Davis, C.A., Zhang, Y., Li, R., Salit, M., Gingeras, T.R., Oliver, B.: Synthetic spike-in standards for RNA-seq experiments. Genome research 21(9), 1543–1551 (2011).

22. Alexander, D.L.J., Tropsha, A., Winkler, D.A.: Beware of R2: simple, unambiguous assessment of the prediction accuracy of QSAR and QSPR models. Journal of chemical information and modeling 55(7), 1316–1322 (2015).

23. Ryvkin, P., Leung, Y.Y., Silverman, I.M., Childress, M., Valladares, O., Dragomir, I., Gregory, B.D., Wang, L.-S.: HAMR: high-throughput annotation of modified ribonucleotides. RNA 19(12), 1684–1692 (2013)

24. Clark, W.C., Evans, M.E., Dominissini, D., Zheng, G., Pan, T.: tRNA base methylation identification and quantification via high-throughput sequencing. RNA (2016).

25. Katibah, G.E., Qin, Y., Sidote, D.J., Yao, J., Lambowitz, A.M., Collins, K.: Broad and adaptable RNA structure recognition by the human interferon-induced tetratricopeptide repeat protein IFIT5. Proceedings of the National Academy of Sciences of the United States of America 111 (33), 12025–12030 (2014)

26. Robert, C., Watson, M.: Errors in RNA-Seq quantification affect genes of relevance to human disease. Genome Biology 16, 177 (2015).

27. Hrdlickova, B., de Almeida, R.C., Borek, Z., Withoff, S.: Genetic variation in the non-coding genome: Involvement of micro-RNAs and long non-coding RNAs in disease. Biochimica et Biophysica Acta (BBA) - Molecular Basis of Disease 1842(10), 1910–1922 (2014).

28. Cunningham, F., Amode, M.R., Barrell, D., Beal, K., Billis, K., Brent, S., Carvalho-Silva, D., Clapham, P., Coates, G., Fitzgerald, S., Gil, L., Girón, C.G., Gordon, L., Hourlier, T., Hunt, S.E., Janacek, S.H., Johnson, N., Juettemann, T., Kähäri, A.K., Keenan, S., Martin, F.J., Maurel, T., McLaren, W., Murphy, D.N., Nag, R., Overduin, B., Parker, A., Patricio, M., Perry, E., Pignatelli, M., Riat, H.S., Sheppard, D., Taylor, K., Thormann, A., Vullo, A., Wilder, S.P., Zadissa, A., Aken, B.L., Birney, E., Harrow, J., Kinsella, R., Muffato, M., Ruffier, M., Searle, S.M.J., Spudich, G., Trevanion, S.J., Yates, A., Zerbino, D.R., Flicek, P.: Ensembl 2015. Nucleic Acids Research 43(D1), 662–669 (2015).

29. Chan, P.P., Lowe, T.M.: GtRNAdb 2.0: an expanded database of transfer RNA genes identified in complete and draft genomes. Nucleic Acids Research 44(D1), 184–189 (2016).

30. Durinck, S., Spellman, P.T., Birney, E., Huber, W.: Mapping Identifiers for the Integration of Genomic Datasets with the R/Bioconductor package biomaRt. Nature protocols 4(8), 1184–1191 (2009).

31. Durinck, S., Moreau, Y., Kasprzyk, A., Davis, S., De Moor, B., Brazma, A., Huber, W.: BioMart and Bioconductor: a powerful link between biological databases and microarray data analysis. Bioinformatics 21(16), 3439–3440 (2005).

32. Martin, M.: Cutadapt removes adapter sequences from high-throughput sequencing reads. EMBnet.journal 17(1), 10–12 (2011).

33. Quinlan, A.R.: BEDTools: the swiss-army tool for genome feature analysis. Current protocols in bioinformatics 47, 11–12134 (2014).

34. Love, M.I., Huber, W., Anders, S.: Moderated estimation of fold change and dispersion for RNA-seq data with DESeq2. Genome Biology 15 (12), 550 (2014).

35. Soneson, C., Love, M., Robinson, M.: differential analyses for RNA-seq: transcript-level estimates improve gene-level inferences [version 1; referees: 2 approved]. F1000Research 4(1521) (2015).

36. Kuhn, M.: Building Predictive Models in R Using the caret Package. Journal of Statistical Software 28(5), 1–26 (2008)

37. Freedman, J.E., Gerstein, M., Mick, E., Rozowsky, J., Levy, D., Kitchen, R., Das, S., Shah, R., Danielson, K., Beaulieu, L., Navarro, F.C.P., Wang, Y., Galeev, T.R., Holman, A., Kwong, R.Y., Murthy, V., Tanriverdi, S.E., Koupenova, M., Mikhalev, E., Tanriverdi, K.: Diverse human extracellular RNAs are widely detected in human plasma. Nature Communications 7, 11106 (2016).

38. Hopper, A.K., Phizicky, E.M.: tRNA transfers to the limelight. Genes & Development 17(2), 162–180 (2003).

